## Supplemental Files Bourgade et al. for "Development of a CRISPR activation system for targeted gene upregulation in *Synechocystis* sp. PCC 6803"

**Bourgade et al, 2024**

Supplementary Figure 1. Tool stability	12

Supplementary Figure 2. Impact of rhamnose concentration on activation levels	12

Supplementary Figure 3. Growth profiles and IB/3M1B production of CRISPRa-activated strains with multiple *kivD*^S286T^ copies	13

Supplementary Figure 4. Growth profiles of strains with CRISPRa targeting for target mapping	14

Supplementary Figure 5. IB/3M1B ratio for CRISPRa-targeted strains	14

Supplementary Figure 6. Relative transcript levels of target genes	15

Supplementary Figure 7. Simplified metabolic map of IB/3M1B biosynthesis	16

Supplementary Figure 8. Relative transcript levels of multiplexed gene targets	17

### Tables

**Supplementary Table 1.** Synechocystis background strains used in this study.

| **Strain** | **Genotype** | **Reference** |
| --- | --- | --- |
| WT | Wild-type *Synechocystis* sp. PCC 6803 | ^1^ |
| sBB_CA1 | Δ*slr0168*::P*_trc_*-*GFP* (Sp^R^*) | This study |
| sBB_CA2 | Δ*slr0168*::J23119-*GFP* (Sp^R^*) | This study |
| sBB_CA3 | Δ*slr0168*::J23116-*GFP* (Sp^R^*) | This study |
| sBB_CA4 | Δ*slr0168*::J23107-*GFP* (Sp^R^*) | This study |
| sBB_CA5 | Δ*slr0168*::J23101-*GFP* (Sp^R^*) | This study |
| ddh_kivD | Δ*ddh*::P*_trc_*-*kivd^S286T^* (Cm^R^*) | Obtained with plasmid pHX8^2^ |
| HX11 | Δ*ddh*::P*_trc_*-*kivd^S286T^* (Cm^R^*), *Δslr0168*:: P*_trc_*-*kivd^S286T^* (Sp^R^*) | ^3^ |
| HX51 | Δ*ddh*::P*_trc_*-*kivd^S286T^* (Cm^R^*); *Δslr0168*:: P*_trc_*-*kivd^S286T^* (Sp^R^*); ); *Δsll1564*:: P*_trc_*-*kivd^S286T^* (Em^R^*) | Xie and Lindblad, unpublished |

*Sp^R^: spectinomycin resistance cassette; Cm^R^: chloramphenicol resistance cassette; Em^R^: erythromycin resistance cassette

3.	Xie, H., Bourgade, B., Stensjö, K. & Lindblad, P. dCas12a-mediated CRISPR interference for multiplex gene repression in cyanobacteria for enhanced isobutanol and 3-methyl-1-butanol production. *Bioresour. Technol.* Under revision.

**Supplementary Table 2.** List of plasmids used in this study.

| **Plasmid** | **Relevant elements** | **Reference** |
| --- | --- | --- |
| P3 | *HA_slr0168_*; P*_trc_*-*sll1363*-*slr0452*; SpR* | ^1^ |
| pBB_CA1 | HA*_slr0168_*; P*_trc_*-GFP; SpR* | This study |
| pBB_CA2 | HA*_slr0168_*; J23119-GFP; SpR* | This study |
| pBB_CA3 | HA*_slr0168_*; J23116-GFP; SpR* | This study |
| pBB_CA4 | HA*_slr0168_*; J23107-GFP; SpR* | This study |
| pBB_CA5 | HA*_slr0168_*; J23101-GFP; SpR* | This study |
| pBB_dCas12a | *rhaS*; P*_rha_*-dCas12; P*_rha_*-gRNA_(no target)_; KmR* | ^2^ |
| pBB_CA | *rhaS*; P*_rha_*-dCas12-SoxS^R93A^; P*_rha_*-_gRNA(no target)_; KmR* | This study |
| pBB_CA6 | *rhaS*; P*_rha_*-dCas12-SoxS^R93A^; P*_rha_*-gRNA_(-48)_; KmR* | This study |
| pBB_CA7 | *rhaS*; P*_rha_*-dCas12-SoxS^R93A^; P*_rha_*-gRNA_(-97 NTS)_; KmR* | This study |
| pBB_CA8 | *rhaS*; P*_rha_*-dCas12-SoxS^R93A^; P*_rha_*-gRNA_(-108)_; KmR* | This study |
| pBB_CA9 | *rhaS*; P*_rha_*-dCas12-SoxS^R93A^; P*_rha_*-gRNA_(-144 NTS)_; KmR* | This study |
| pBB_CA10 | *rhaS*; P*_rha_*-dCas12-SoxS^R93A^; P*_rha_*-gRNA_(-156)_; KmR* | This study |
| pBB_CA11 | *rhaS*; P*_rha_*-dCas12-SoxS^R93A^; P*_rha_*-gRNA_(-251)_; KmR* | This study |
| pBB_CA12 | *rhaS*; P*_rha_*-dCas12-SoxS^R93A^; P*_rha_*-gRNA_(-328)_; KmR* | This study |
| pBB_CA13 | *rhaS*; P*_rha_*-dCas12-SoxS^R93A^; P*_rha_*-gRNA_(-108/-156)_; KmR* | This study |
| pBB_CA14 | *rhaS*; P*_rha_*-dCas12-SoxS^R93A^; P*_rha_*-gRNA_(CDS)_; KmR* | This study |
| pBB_CA15 | *rhaS*; P*_rha_*-dCas12-SoxS^R93A^; P*_rha_*-gRNA_(J23)_; KmR* | This study |
| pBB_CA16 | *rhaS*; P*_rha_*-dCas12-SoxS^R93A^; P*_rha_*-gRNA_(_*_ddh_*_)_; KmR* | This study |
| pBB_CA17 | *rhaS*; P*_rha_*-dCas12-SoxS^R93A^; P*_rha_*-gRNA_(_*_NS1_*_)_; KmR* | This study |
| pBB_CA18 | *rhaS*; P*_rha_*-dCas12-SoxS^R93A^; P*_rha_*-gRNA_(_*_NS1_*_-_*_ddh_*_)_; KmR* | This study |
| pBB_CA19 | *rhaS*; P*_rha_*-dCas12-SoxS^R93A^; P*_rha_*-gRNA_(_*_sll1654_*_)_; KmR* | This study |
| pBB_CA20 | *rhaS*; P*_rha_*-dCas12-SoxS^R93A^; P*_rha_*-gRNA_(_*_NS1_*_-_*_sll1564_*_)_; KmR* | This study |
| pBB_CA21 | *rhaS*; P*_rha_*-dCas12-SoxS^R93A^; P*_rha_*-gRNA_(_*_ddh_*_-_*_sll1564_*_)_; KmR* | This study |
| pBB_CA22 | *rhaS*; P*_rha_*-dCas12-SoxS^R93A^; P*_rha_*-gRNA_(Triple)_; KmR* | This study |
| pBB_CA23 | *rhaS*; P*_rha_*-dCas12-SoxS^R93A^; P*_rha_*-gRNA_(_*_pyk2_*_.1)_; KmR* | This study |
| pBB_CA24 | *rhaS*; P*_rha_*-dCas12-SoxS^R93A^; P*_rha_*-gRNA_(_*_pyk2_*_.2)_; KmR* | This study |
| pBB_CA25 | *rhaS*; P*_rha_*-dCas12-SoxS^R93A^; P*_rha_*-gRNA_(_*_pyk1_*_.1)_; KmR* | This study |
| pBB_CA26 | *rhaS*; P*_rha_*-dCas12-SoxS^R93A^; P*_rha_*-gRNA_(_*_pyk1_*_.2)_; KmR* | This study |
| pBB_CA27 | *rhaS*; P*_rha_*-dCas12-SoxS^R93A^; P*_rha_*-gRNA_(_*_pntA_*_.1)_; KmR* | This study |
| pBB_CA28 | *rhaS*; P*_rha_*-dCas12-SoxS^R93A^; P*_rha_*-gRNA_(_*_pntA_*_.2)_; KmR* | This study |
| pBB_CA29 | *rhaS*; P*_rha_*-dCas12-SoxS^R93A^; P*_rha_*-gRNA_(ME)_; KmR* | This study |
| pBB_CA30 | *rhaS*; P*_rha_*-dCas12-SoxS^R93A^; P*_rha_*-gRNA_(tpi)_; KmR* | This study |
| pBB_CA31 | *rhaS*; P*_rha_*-dCas12-SoxS^R93A^; P*_rha_*-gRNA_(_*_petH_*_)_; KmR* | This study |
| pBB_CA32 | *rhaS*; P*_rha_*-dCas12-SoxS^R93A^; P*_rha_*-gRNA_(_*_acnSP_*_)_; KmR* | This study |
| pBB_CA33 | *rhaS*; P*_rha_*-dCas12-SoxS^R93A^; P*_rha_*-gRNA_(_*_slr6040_*_)_; KmR* | This study |
| pBB_CA34 | *rhaS*; P*_rha_*-dCas12-SoxS^R93A^; P*_rha_*-gRNA_(_*_pyk2_*_.1-_ *_pyk2_*_.2)_; KmR* | This study |
| pBB_CA35 | *rhaS*; P*_rha_*-dCas12-SoxS^R93A^; P*_rha_*-gRNA_(_*_pyk1_*_.1-_ *_pyk1_*_.2)_; KmR* | This study |
| pBB_CA36 | *rhaS*; P*_rha_*-dCas12-SoxS^R93A^; P*_rha_*-gRNA_(_*_pyk2_*_.1-_ *_pyk1_*_.1)_; KmR* | This study |
| pBB_CA37 | *rhaS*; P*_rha_*-dCas12-SoxS^R93A^; P*_rha_*-gRNA_(_*_pyk2_*_.1-ME)_; KmR* | This study |
| pBB_CA38 | *rhaS*; P*_rha_*-dCas12-SoxS^R93A^; P*_rha_*-gRNA_(_*_acnSP_*_-ME)_; KmR* | This study |
| pBB_CA39 | *rhaS*; P*_rha_*-dCas12-SoxS^R93A^; P*_rha_*-gRNA_(_*_pyk2_*_.1-_ *_slr6040_*_)_; KmR* | This study |
| pBB_CA40 | *rhaS*; P*_rha_*-dCas12-SoxS^R93A^; P*_rha_*-gRNA_(_*_slr6040_*_-ME)_; KmR* | This study |

*Sp^R^: spectinomycin resistance cassette; Km^R^: kanamycin resistance cassette

2.	Xie, H., Bourgade, B., Stensjö, K. & Lindblad, P. dCas12a-mediated CRISPR interference for multiplex gene repression in cyanobacteria for enhanced isobutanol and 3-methyl-1-butanol production. *Bioresour. Technol.* Under revision.

Supplementary Table 3. Oligonucleotides used in this study.

| I. **Primers for building plasmids** | |
| --- | --- |
| Backbone_Fwd | AGATCTAACTACCGCATTAAAG |
| Backbone_Rev | GATTATCGGCACCGTCTCTAATTTTAACGTGGCTTTGCGC |
| dCas12_Fwd | TTAATGCGGTAGTTAGATCTTTGACAGCTAGCTCAGTCC |
| dCas12_Rev | AGCGCGTCCAGATCCAGAAGCCTCAGATCCGTTATTCCTATTCTGCACGA |
| SoxS_Fwd | GGATCTGAGGCTTCTGGATCTGGACGCGCTTCCCACCAGAAAATCATCCA |
| SoxS_Rev | TTAGAGACGGTGCCGATAATCTTAGAGACGGTGCCGATAATC |
| Ptrc_Fwd | ATATATGAATTCGAGCTGTTGACAATTGTGAG |
| Ptrc_Rev | GAAAAGTTCTTCTCCTTTACTCATCATTAGAAAACCTCCTTAGC |
| GFP_Fwd | TCATGCTAAGGAGGTTTTCTAATGATGAGTAAAGGAGAAGAACT |
| GFP_Rev | ATATATGCGGCCGCTTATTTGTATAGTTCATCCA |
| J23119_Fwd | ATATATGAATTCTTGACAGCTAGCTCAGTCCTAGGTATAATGCTAGCGGGCCCAAGTTCACTT |
| J23116_Fwd | ATATATGAATTCTTGACAGCTAGCTCAGTCCTAGGGACTATGCTAGCTGGGCCCAAGTTCACTT |
| J23107_Fwd | ATATATGAATTCTTTACGGCTAGCTCAGCCCTAGGTATTATGCTAGCTGGGCCCAAGTTCACTT |
| J23101_Fwd | ATATATGAATTCTTTACAGCTAGCTCAGTCCTAGGTATTATGCTAGCTGGGCCCAAGTTCACTT |
| II. **Primers for cloning gRNAs** | |
| -48_Fwd | 5’Phos/ AGATAATTCGAGCTGTTGACAATT |
| -48_Rev | 5’Phos/ AGACAATTGTCAACAGCTCGAATT |
| -97_Fwd | 5’Phos/ AGATATTTTAGATTAATTCAACAG |
| -97_Rev | 5’Phos/ AGACCTGTTGAATTAATCTAAAAT |
| -108_Fwd | 5’Phos/ AGATTGAAATATTACTGTTGAATT |
| -108_Rev | 5’Phos/ AGACAATTCAACAGTAATATTTCA |
| -144_Fwd | 5’Phos/ AGATAACTCGCAATAATTGCATTA |
| -144_Rev | 5’Phos/ AGACTAATGCAATTATTGCGAGTT |
| -156_Fwd | 5’Phos/ AGATAATCAACTTAATTAATGCAA |
| -156_Rev | 5’Phos/ AGACTTGCATTAATTAAGTTGATT |
| -251_Fwd | 5’Phos/ AGATATTGAAGAAATGGCCCTGGA |
| -251_Rev | 5’Phos/ AGACTCCAGGGCCATTTCTTCAAT |
| -328_Fwd | 5’Phos/ AGATTTCATTGTGTTAGGGGAGGT |
| -328_Rev | 5’Phos/ AGACACCTCCCCTAACACAATGAA |
| CDS_Fwd | 5’Phos/ AGATTCTTATGGTGTTCAATGCTT |
| CDS_Rev | 5’Phos/ AGACAAGCATTGAACACCATAAGA |
| -108/-156_Fwd | 5’Phos/ AGATAATCAACTTAATTAATGCAAGTCTAAGAACTTTAAATAATTTCTACTGTTGTAGATTGAAATATTACTGTTGAATT |
| -108/-156_Rev | 5’Phos/ AGACAATTCAACAGTAATATTTCAATCTACAACAGTAGAAATTATTTAAAGTTCTTAGACTTGCATTAATTAAGTTGATT |
| ddh_Fwd | 5’Phos/ AGATCAAACACGTTCTAAACTACT |
| ddh_Rev | 5’Phos/ AGACAGTAGTTTAGAACGTGTTTG |
| NS1_Fwd | 5’Phos/ AGATAATCAACTTAATTAATGCAA |
| NS1_Rev | 5’Phos/ AGACTTGCATTAATTAAGTTGATT |
| sll1564_Fwd | 5’Phos/ AGATAAATCCAGTAACTACATAAT |
| sll1564_Rev | 5’Phos/ AGACATTATGTAGTTACTGGATTT |
| NS1-ddh_Fwd | 5’Phos/ AGATCAAACACGTTCTAAACTACTGTCTAAGAACTTTAAATAATTTCTACTGTTGTAGATAATCAACTTAATTAATGCAA |
| NS1-ddh_Rev | 5’Phos/ AGACTTGCATTAATTAAGTTGATTATCTACAACAGTAGAAATTATTTAAAGTTCTTAGACAGTAGTTTAGAACGTGTTTG |
| NS1-sll1564_Fwd | 5’Phos/ AGATAATCAACTTAATTAATGCAAGTCTAAGAACTTTAAATAATTTCTACTGTTGTAGATAAATCCAGTAACTACATAAT |
| NS1-sll1564_Rev | 5’Phos/ AGACATTATGTAGTTACTGGATTTATCTACAACAGTAGAAATTATTTAAAGTTCTTAGACTTGCATTAATTAAGTTGATT |
| ddh-sll1564_Fwd | 5’Phos/ AGATCAAACACGTTCTAAACTACTGTCTAAGAACTTTAAATAATTTCTACTGTTGTAGATAAATCCAGTAACTACATAAT |
| ddh-sll1564_Rev | 5’Phos/ AGACATTATGTAGTTACTGGATTTATCTACAACAGTAGAAATTATTTAAAGTTCTTAGACAGTAGTTTAGAACGTGTTTG |
| Triple1_Fwd | 5’Phos/ AGATCAAACACGTTCTAAACTACTGTCTAAGAACTTTAAATAATTTCTACTGTTGTAGATAATCAACTTAATTAATGCAA |
| Triple1_Rev | ATCTACAACAGTAGAAATTATTTAAAGTTCTTAGACAGTAGTTTAGAACGTGTTTG |
| Triple2_Fwd | GTCTAAGAACTTTAAATAATTTCTACTGTTGTAGATAAATCCAGTAACTACATAAT |
| Triple2_Rev | 5’Phos/ AGACATTATGTAGTTACTGGATTTATCTACAACAGTAGAAATTATTTAAAGTTCTTAGACTTGCATTAATTAAGTTGATT |
| pyk2.1_Fwd | 5’Phos/ AGATAGTTGCATCGGCTTACAGGG |
| pyk2.1_Rev | 5’Phos/ AGACCCCTGTAAGCCGATGCAACT |
| pyk2.2_Fwd | 5’Phos/ AGATGGCAATTTTTCCCAATAGTC |
| pyk2.2_Rev | 5’Phos/ AGACGACTATTGGGAAAAATTGCC |
| pyk1.1_Fwd | 5’Phos/ AGATAAAATTAAAACCGCCGGTAA |
| pyk1.1_Rev | 5’Phos/ AGACTTACCGGCGGTTTTAATTTT |
| pyk1.2_Fwd | 5’Phos/ AGATCCCGGCAAAATATAATCCAG |
| pyk1.2_Rev | 5’Phos/ AGACCTGGATTATATTTTGCCGGG |
| pntA.1_Fwd | 5’Phos/ AGATCGCTGTCCCCGTAAGATGGG |
| pntA.1_Rev | 5’Phos/ AGACCCCATCTTACGGGGACAGCG |
| pntA.2_Fwd | 5’Phos/ AGATCTCACAAATTAGCCTTAACG |
| pntA.2_Rev | 5’Phos/ AGACCGTTAAGGCTAATTTGTGAG |
| ME_Fwd | 5’Phos/ AGATCCCATGTATCTAGCCCCATC |
| ME_Rev | 5’Phos/ AGACGATGGGGCTAGATACATGGG |
| Tpi_Fwd | 5’Phos/ AGATAATCTAGTCCAATCGAGAAG |
| Tpi_Rev | 5’Phos/ AGACCTTCTCGATTGGACTAGATT |
| petH_Fwd | 5’Phos/ AGATCCGGACGATTAACCCTGGAA |
| petH_Rev | 5’Phos/ AGACTTCCAGGGTTAATCGTCCGG |
| pyk2.1.2_Fwd | 5’Phos/ AGATAGTTGCATCGGCTTACAGGGGTCTAAGAACTTTAAATAATTTCTACTGTTGTAGATGGCAATTTTTCCCAATAGTC |
| pyk2.1.2_Rev | 5’Phos/ AGACGACTATTGGGAAAAATTGCCATCTACAACAGTAGAAATTATTTAAAGTTCTTAGACCCCTGTAAGCCGATGCAACT |
| pyk1.1.2_Fwd | 5’Phos/ AGATAAAATTAAAACCGCCGGTAAGTCTAAGAACTTTAAATAATTTCTACTGTTGTAGATCCCGGCAAAATATAATCCAG |
| pyk1.1.2_Rev | 5’Phos/ AGACCTGGATTATATTTTGCCGGGATCTACAACAGTAGAAATTATTTAAAGTTCTTAGACTTACCGGCGGTTTTAATTTT |
| pyk2.1-pyk1.1_Fwd | 5’Phos/ AGATAGTTGCATCGGCTTACAGGGGTCTAAGAACTTTAAATAATTTCTACTGTTGTAGATAAAATTAAAACCGCCGGTAA |
| pyk2.1-pyk1.1_Rev | 5’Phos/ AGACTTACCGGCGGTTTTAATTTTATCTACAACAGTAGAAATTATTTAAAGTTCTTAGACCCCTGTAAGCCGATGCAACT |
| pyk2.1-ME_Fwd | 5’Phos/ AGATAGTTGCATCGGCTTACAGGGGTCTAAGAACTTTAAATAATTTCTACTGTTGTAGATCCCATGTATCTAGCCCCATC |
| pyk2.1-ME_Rev | 5’Phos/ AGACGATGGGGCTAGATACATGGGATCTACAACAGTAGAAATTATTTAAAGTTCTTAGACCCCTGTAAGCCGATGCAACT |
| slr6040_Fwd | 5’Phos/ AGATGACCAGGGTTGGGTGAACTA |
| slr6040_Rev | 5’Phos/ AGACTAGTTCACCCAACCCTGGTC |
| slr6040-ME_Fwd | 5’Phos/ AGATGACCAGGGTTGGGTGAACTAGTCTAAGAACTTTAAATAATTTCTACTGTTGTAGATCCCATGTATCTAGCCCCATC |
| slr6040-ME_Rev | 5’Phos/ AGACGATGGGGCTAGATACATGGGATCTACAACAGTAGAAATTATTTAAAGTTCTTAGACTAGTTCACCCAACCCTGGTC |
| slr6040-pyk2.1_Fwd | 5’Phos/ AGATGACCAGGGTTGGGTGAACTAGTCTAAGAACTTTAAATAATTTCTACTGTTGTAGATAGTTGCATCGGCTTACAGGG |
| slr6040- pyk2.1_Rev | 5’Phos/ AGACCCCTGTAAGCCGATGCAACTATCTACAACAGTAGAAATTATTTAAAGTTCTTAGACTAGTTCACCCAACCCTGGTC |
| acnSP_Fwd | 5’Phos/ AGATGCGATTGGCAGCGTGGCGAC |
| acnSP_Rev | 5’Phos/ AGACGTCGCCACGCTGCCAATCGC |
| acnSP-ME_Fwd | AGATGCGATTGGCAGCGTGGCGACGTCTAAGAACTTTAAATAATTTCTACTGTTGTAGATCCCATGTATCTAGCCCCATC |
| acnSP-ME_Rev | AGACGATGGGGCTAGATACATGGGATCTACAACAGTAGAAATTATTTAAAGTTCTTAGACGTCGCCACGCTGCCAATCGC |
| III. **Primers for RT-qPCR** | |
| q.rnpB_Fwd | CGTTAGGATAGTGCCACAG |
| q.rnpB_Rev | CGCTCTTACCGCACCTTTG |
| q.GFP_Fwd | GATGGAAGCGTTCAACTAGCA |
| q.GFP_Rev | GCAGATTGTGTGGACAGGTAAT |
| q.KivD_Fwd | GAGTGAACCCAACCTGAAAGA |
| q.KivD_Rev | TGGGTAAAGGCTCCAGTAGA |
| q.Flag_Fwd | GACTACAAGGATGACGATGACAA |
| q.His_Fwd | CATCACCATCACCACGGTAG |
| q.Tag_Rev | GTTCATGTAAGCGGTCCAGTAA |
| q.pyk2_Fwd | TCCAGCCCGAACATCAAATC |
| q.pyk2_Rev | GGGAACCCTTTCCAGCAATAA |
| q.pyk1_Fwd | TGGAAGCGAGGAGAGAAGTA |
| q.pyk1_Rev | AACCCATGCCCTAAGTGAAG |
| q.pntA_Fwd | CCAGTCAATCCAGTCAGCTTTA |
| q.pntA_Rev | CCTCATCTTCCACATCCACTTT |
| q.ME_Fwd | GGCAGTGAAAGCTCTGGATAA |
| q.ME_Rev | AATGCGACTAACCACGCTAAT |
| q.tpi_Fwd | CAATCTAACCTAGTCATCGCCTAC |
| q.tpi_Rev | TCCCGAATCAGCCCAATAAC |
| q.petH_Fwd | CCGTTACCTAGAAGGGCAAAG |
| q.petH_Rev | GTCTGGTGGAAGCAATGGAATA |
| q.slr6040_Fwd | GATCTAGGAATGGTGCTGGAAA |
| q.slr6040_Rev | CACCCAACCCTGGTCTAAAT |
| q.acnSP_Fwd | CGTCGCCACGCTGCCAATCG |
| q.acnSP_Rev | GGATTTTTAGCAGTTCACAT |

**Supplementary Table 4.** DNA sequences of relevant CRISPRa elements used in this study.

| **Element** | **DNA sequence** | **Reference** |
| --- | --- | --- |
| rhaS | ATGACCGTATTACATAGTGTGGATTTTTTTCCGTCTGGTAACGCGTCCGTGGCGATAGAACCCCGGCTCCCGCAGGCGGATTTTCCTGAACATCATCATGATTTTCATGAAATTGTGATTGTCGAACATGGCACGGGTATTCATGTGTTTAATGGGCAGCCCTATACCATCACCGGTGGCACGGTCTGTTTCGTACGCGATCATGATCGGCATCTGTATGAACATACCGATAATCTGTGTCTGACCAATGTGCTGTATCGCTCGCCGGATCGATTTCAGTTTCTCGCCGGGCTGAATCAGTTGCTGCCACAAGAGCTGGATGGGCAGTATCCGTCTCACTGGCGCGTTAACCACAGCGTCTTGCAGCAAGTGCGACAGCTGGTTGCACAGATGGAACAGCAGGAAGGGGAAAATGATTTACCCTCGACCGCCAGTCGCGAGATCTTGTTTATGCAATTACTGCTCTTGCTGCGTAAAAGCAGTTTGCAGGAGAACCTGGAAAACAGCGCATCACGTCTCAACTTGCTTCTGGCCTGGCTGGAGGACCATTTTGCCGATGAGGTGAATTGGGATGCCGTGGCGGATCAATTTTCTCTTTCACTGCGTACGCTACATCGGCAGCTTAAGCAGCAAACGGGACTGACGCCTCAGCTATACCTGAACCGCCTGCGATTGATGAAAGCCCGACATCTGCTACGCCACAGCGAGGCCAGCGTTACTGACATCGCCTATCGCTGTGGATTCAGCGACAGTAACCACTTTTCGACGCTTTTTCGCCGAGAGTTTAACTGGTCACCGCGTGATATTCGCCAGGGACGGGATGGCTTTCTGCAATAA | ^1^ |
| P*_rha_* | GCCACAATTCAGCAAATTGTGAACATCATCACGTTCATCTTTCCCTGGTTGCCAATGGCCCATTTTCCTGTCAGTAACGAGAAGGTCGCGAATTCAGGCGCTTTTTAGACTGGTCGTAATGAA | ^2^ |
| dCas12a | ATGTCAATTTATCAAGAATTTGTTAATAAATATAGTTTAAGTAAAACTCTAAGATTTGAGTTAATCCCACAGGGTAAAACACTTGAAAACATAAAAGCAAGAGGTTTGATTTTAGATGATGAGAAAAGAGCTAAAGACTACAAAAAGGCTAAACAAATAATTGATAAATATCATCAGTTTTTTATAGAGGAGATATTAAGTTCGGTTTGTATTAGCGAAGATTTATTACAAAACTATTCTGATGTTTATTTTAAACTTAAAAAGAGTGATGATGATAATCTACAAAAAGATTTTAAAAGTGCAAAAGATACGATAAAGAAACAAATATCTGAATATATAAAGGACTCAGAGAAATTTAAGAATTTGTTTAATCAAAACCTTATCGATGCTAAAAAAGGGCAAGAGTCAGATTTAATTCTATGGCTAAAGCAATCTAAGGATAATGGTATAGAACTATTTAAAGCCAATAGTGATATCACAGATATAGATGAGGCGTTAGAAATAATCAAATCTTTTAAAGGTTGGACAACTTATTTTAAGGGTTTTCATGAAAATAGAAAAAATGTTTATAGTAGCAATGATATTCCTACATCTATTATTTATAGGATAGTAGATGATAATTTGCCTAAATTTCTAGAAAATAAAGCTAAGTATGAGAGTTTAAAAGACAAAGCTCCAGAAGCTATAAACTATGAACAAATTAAAAAAGATTTGGCAGAAGAGCTAACCTTTGATATTGACTACAAAACATCTGAAGTTAATCAAAGAGTTTTTTCACTTGATGAAGTTTTTGAGATAGCAAACTTTAATAATTATCTAAATCAAAGTGGTATTACTAAATTTAATACTATTATTGGTGGTAAATTTGTAAATGGTGAAAATACAAAGAGAAAAGGTATAAATGAATATATAAATCTATACTCACAGCAAATAAATGATAAAACACTCAAAAAATATAAAATGAGTGTTTTATTTAAGCAAATTTTAAGTGATACAGAATCTAAATCTTTTGTAATTGATAAGTTAGAAGATGATAGTGATGTAGTTACAACGATGCAAAGTTTTTATGAGCAAATAGCAGCTTTTAAAACAGTAGAAGAAAAATCTATTAAAGAAACACTATCTTTATTATTTGATGATTTAAAAGCTCAAAAACTTGATTTGAGTAAAATTTATTTTAAAAATGATAAATCTCTTACTGATCTATCACAACAAGTTTTTGATGATTATAGTGTTATTGGTACAGCGGTACTAGAATATATAACTCAACAAATAGCACCTAAAAATCTTGATAACCCTAGTAAGAAAGAGCAAGAATTAATAGCCAAAAAAACTGAAAAAGCAAAATACTTATCTCTAGAAACTATAAAGCTTGCCTTAGAAGAATTTAATAAGCATAGAGATATAGATAAACAGTGTAGGTTTGAAGAAATACTTGCAAACTTTGCGGCTATTCCGATGATATTTGATGAAATAGCTCAAAACAAAGACAATTTGGCACAGATATCTATCAAATATCAAAATCAAGGTAAAAAAGACCTACTTCAAGCTAGTGCGGAAGATGATGTTAAAGCTATCAAGGATCTTTTAGATCAAACTAATAATCTCTTACATAAACTAAAAATATTTCATATTAGTCAGTCAGAAGATAAGGCAAATATTTTAGACAAGGATGAGCATTTTTATCTAGTATTTGAGGAGTGCTACTTTGAGCTAGCGAATATAGTGCCTCTTTATAACAAAATTAGAAACTATATAACTCAAAAGCCATATAGTGATGAGAAATTTAAGCTCAATTTTGAGAACTCGACTTTGGCTAATGGTTGGGATAAAAATAAAGAGCCTGACAATACGGCAATTTTATTTATCAAAGATGATAAATATTATCTGGGTGTGATGAATAAGAAAAATAACAAAATATTTGATGATAAAGCTATCAAAGAAAATAAAGGCGAGGGTTATAAAAAAATTGTTTATAAACTTTTACCTGGCGCAAATAAAATGTTACCTAAGGTTTTCTTTTCTGCTAAATCTATAAAATTTTATAATCCTAGTGAAGATATACTTAGAATAAGAAATCATTCCACACATACAAAAAATGGTAGTCCTCAAAAAGGATATGAAAAATTTGAGTTTAATATTGAAGATTGCCGAAAATTTATAGATTTTTATAAACAGTCTATAAGTAAGCATCCGGAGTGGAAAGATTTTGGATTTAGATTTTCTGATACTCAAAGATATAATTCTATAGATGAATTTTATAGAGAAGTTGAAAATCAAGGCTACAAACTAACTTTTGAAAATATATCAGAGAGCTATATTGATAGCGTAGTTAATCAGGGTAAATTGTACCTATTCCAAATCTATAATAAAGATTTTTCAGCTTATAGCAAAGGGCGACCAAATCTACATACTTTATATTGGAAAGCGCTGTTTGATGAGAGAAATCTTCAAGATGTGGTTTATAAGCTAAATGGTGAGGCAGAGCTTTTTTATCGTAAACAATCAATACCTAAAAAAATCACTCACCCAGCTAAAGAGGCAATAGCTAATAAAAACAAAGATAATCCTAAAAAAGAGAGTGTTTTTGAATATGATTTAATCAAAGATAAACGCTTTACTGAAGATAAGTTTTTCTTTCACTGTCCTATTACAATCAATTTTAAATCTAGTGGAGCTAATAAGTTTAATGATGAAATCAATTTATTGCTAAAAGAAAAAGCAAATGATGTTCATATATTAAGTATAGCTAGAGGTGAAAGACATTTAGCTTACTATACTTTGGTAGATGGTAAAGGCAATATCATCAAACAAGATACTTTCAACATCATTGGTAATGATAGAATGAAAACAAACTACCATGATAAGCTTGCTGCAATAGAGAAAGATAGGGATTCAGCTAGGAAAGACTGGAAAAAGATAAATAACATCAAAGAGATGAAAGAGGGCTATCTATCTCAGGTAGTTCATGAAATAGCTAAGCTAGTTATAGAGTATAATGCTATTGTGGTTTTTGAGGATTTAAATTTTGGATTTAAAAGAGGGCGTTTCAAGGTAGAGAAGCAGGTCTATCAAAAGTTAGAAAAAATGCTAATTGAGAAACTAAACTATCTAGTTTTCAAAGATAATGAGTTTGATAAAACTGGGGGAGTGCTTAGAGCTTATCAGCTAACAGCACCTTTTGAGACTTTTAAAAAGATGGGTAAACAAACAGGTATTATCTACTATGTACCAGCTGGTTTTACTTCAAAAATTTGTCCTGTAACTGGTTTTGTAAATCAGTTATATCCTAAGTATGAAAGTGTCAGCAAATCTCAAGAGTTCTTTAGTAAGTTTGACAAGATTTGTTATAACCTTGATAAGGGCTATTTTGAGTTTAGTTTTGATTATAAAAACTTTGGTGACAAGGCTGCCAAAGGCAAGTGGACTATAGCTAGCTTTGGGAGTAGATTGATTAACTTTAGAAATTCAGATAAAAATCATAATTGGGATACTCGAGAAGTTTATCCAACTAAAGAGTTGGAGAAATTGCTAAAAGATTATTCTATCGAATATGGGCATGGCGAATGTATCAAAGCAGCTATTTGCGGTGAGAGCGACAAAAAGTTTTTTGCTAAGCTAACTAGTGTCCTAAATACTATCTTACAAATGCGTAACTCAAAAACAGGTACTGAGTTAGATTATCTAATTTCACCAGTAGCAGATGTAAATGGCAATTTCTTTGATTCGCGACAGGCGCCAAAAAATATGCCTCAAGATGCTGATGCCAATGGTGCTTATCATATTGGGCTAAAAGGTCTGATGCTACTAGGTAGGATCAAAAATAATCAAGAGGGCAAAAAACTCAATTTGGTTATCAAAAATGAAGAGTATTTTGAGTTCGTGCAGAATAGGAATAAC | ^3^ |
| SoxS^R93A^ | TCCCACCAGAAAATCATCCAAGATCTGATTGCATGGATCGACGAGCACATTGACCAGCCTCTCAATATCGATGTAGTTGCAAAGAAGAGCGGATACAGCAAATGGTATCTGCAACGCATGTTCCGCACTGTTACACACCAAACATTGGGAGATTATATTCGCCAACGACGACTCCTGCTGGCAGCCGTAGAGTTACGAACTACAGAGCGTCCTATTTTTGACATTGCTATGGATTTGGGTTATGTGAGCCAGCAGACATTCTCCCGTGTGTTTGCGCGTCAGTTTGACCGTACTCCCTCCGATTATCGGCACCGTCTCTAA | This study |

### Figures

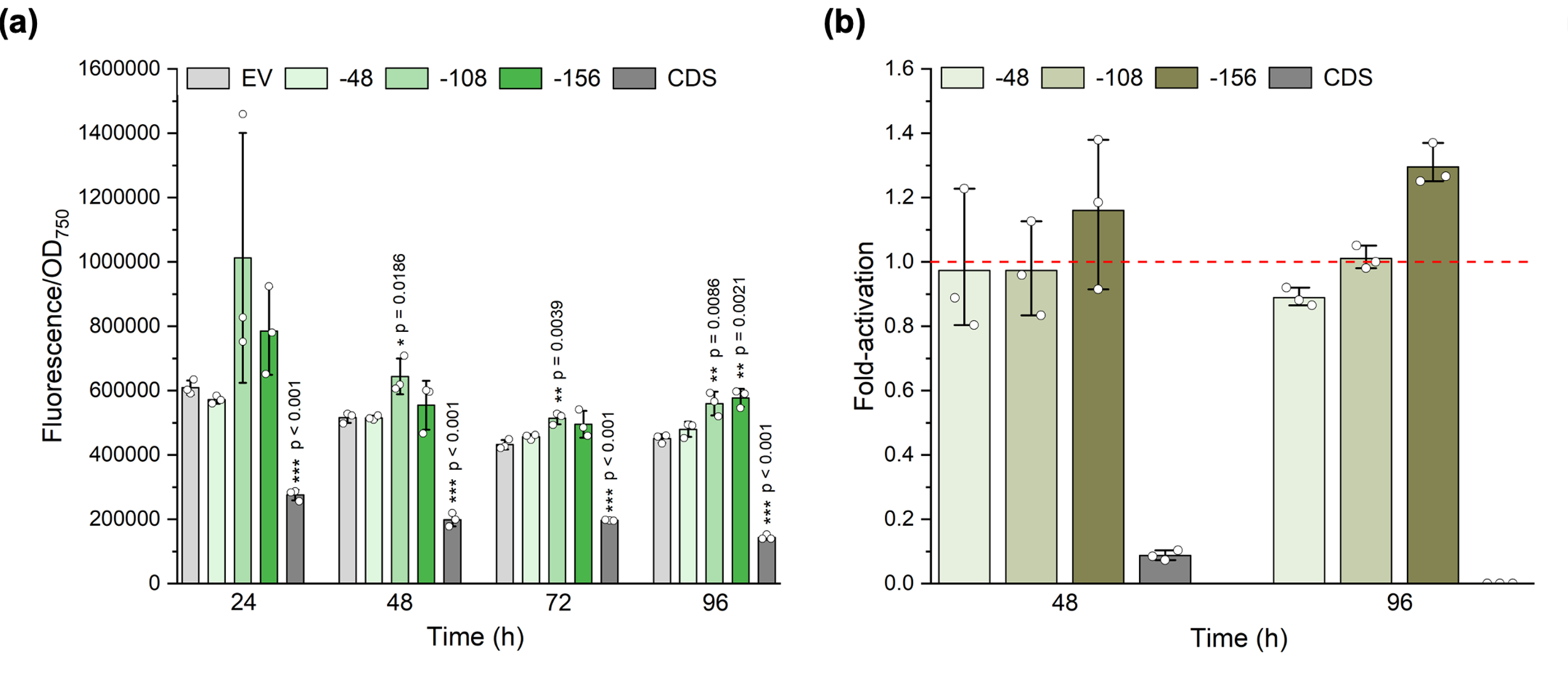

Supplementary Figure 1. GFP activation throughout a 96-hour post-induction period with four selected gRNAs. (a) GFP fluorescence was quantified at 24, 48, 72 and 96 hours after rhamnose induction for four selected gRNAs. (b) Fold-activation was calculated relative to the negative control (EV) at 48h and 96h. EV: negative control; CDS: coding sequence. Error bars indicate standard deviation (n=3). p value representation: * < 0.05; ** < 0.01; *** < 0.001. p value was calculated by comparing each sample to the respective negative control.

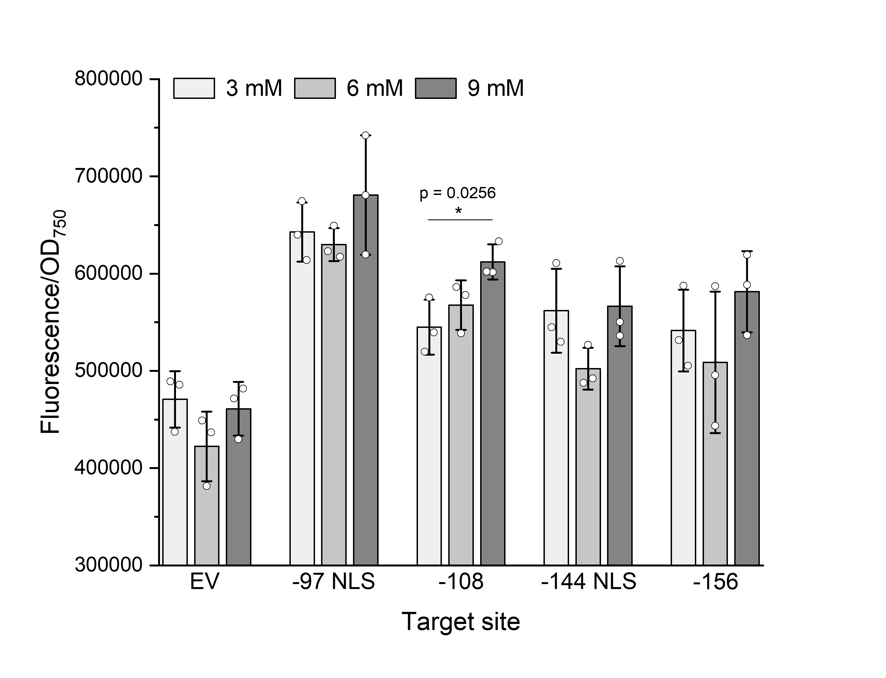

Supplementary Figure 2. Correlation between rhamnose concentration and GFP fluorescence 72 hours post-induction for four selected gRNAs. GFP fluorescence was measured 72 hours post-induction with 3; 6 or 9 mM of rhamnose for four selected gRNAs. Error bars indicate standard deviation (n=3). p value representation: * < 0.05. p value was calculated by comparing each sample to the respective negative control.

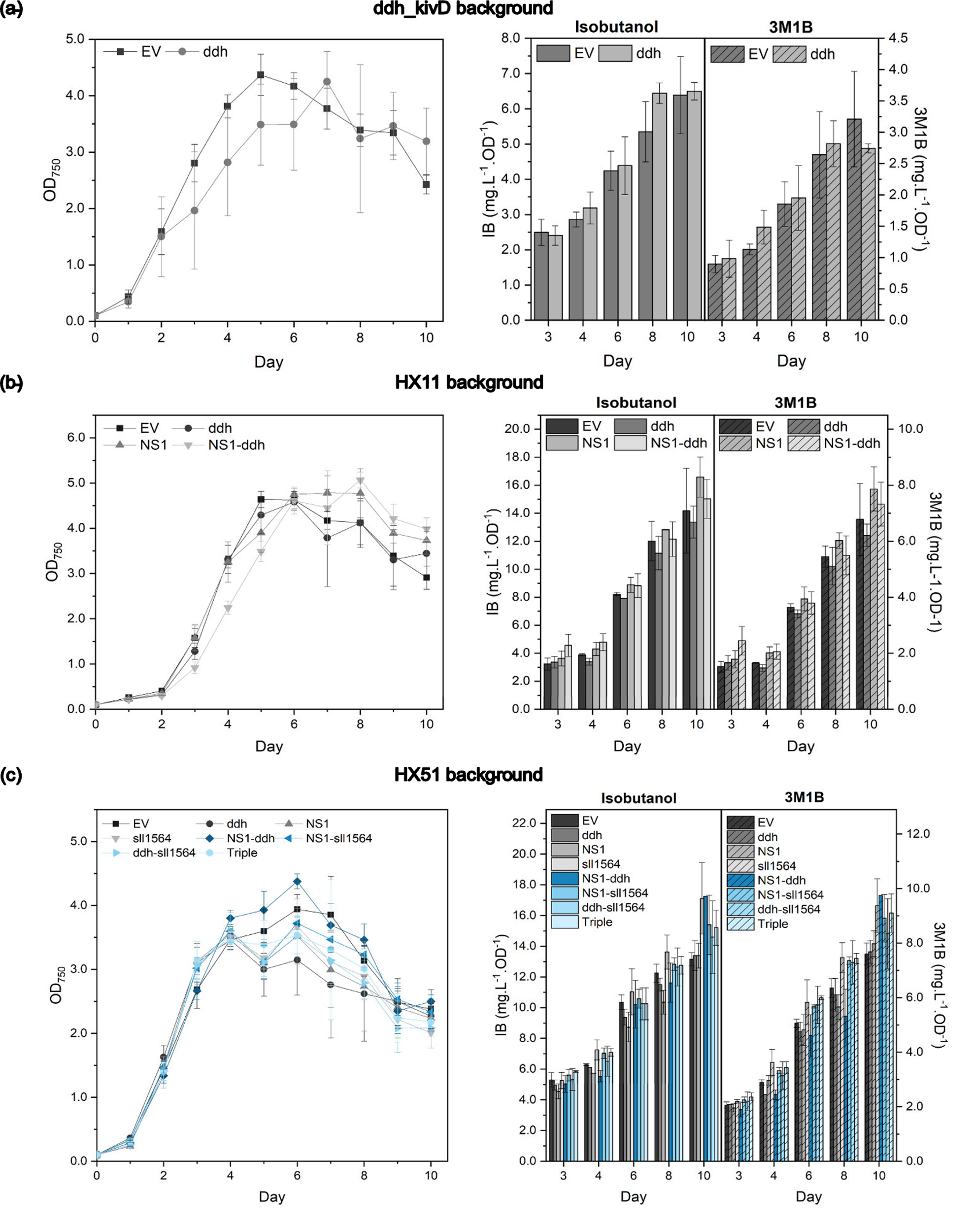

Supplementary Figure 3. Growth profiles and isobutanol (IB)/3-methyl-1-butanol (3M1B) production of CRISPR-activated strains. Growth and corresponding IB/3M1B titres were monitored in CRISPRa-activated derivatives of (a) ddh_kivD, (b) HX11 and (c) HX51 background strains during 10 days post-induction with 3 mM rhamnose**.** EV: negative control. Error bars represent standard deviation (n=3).

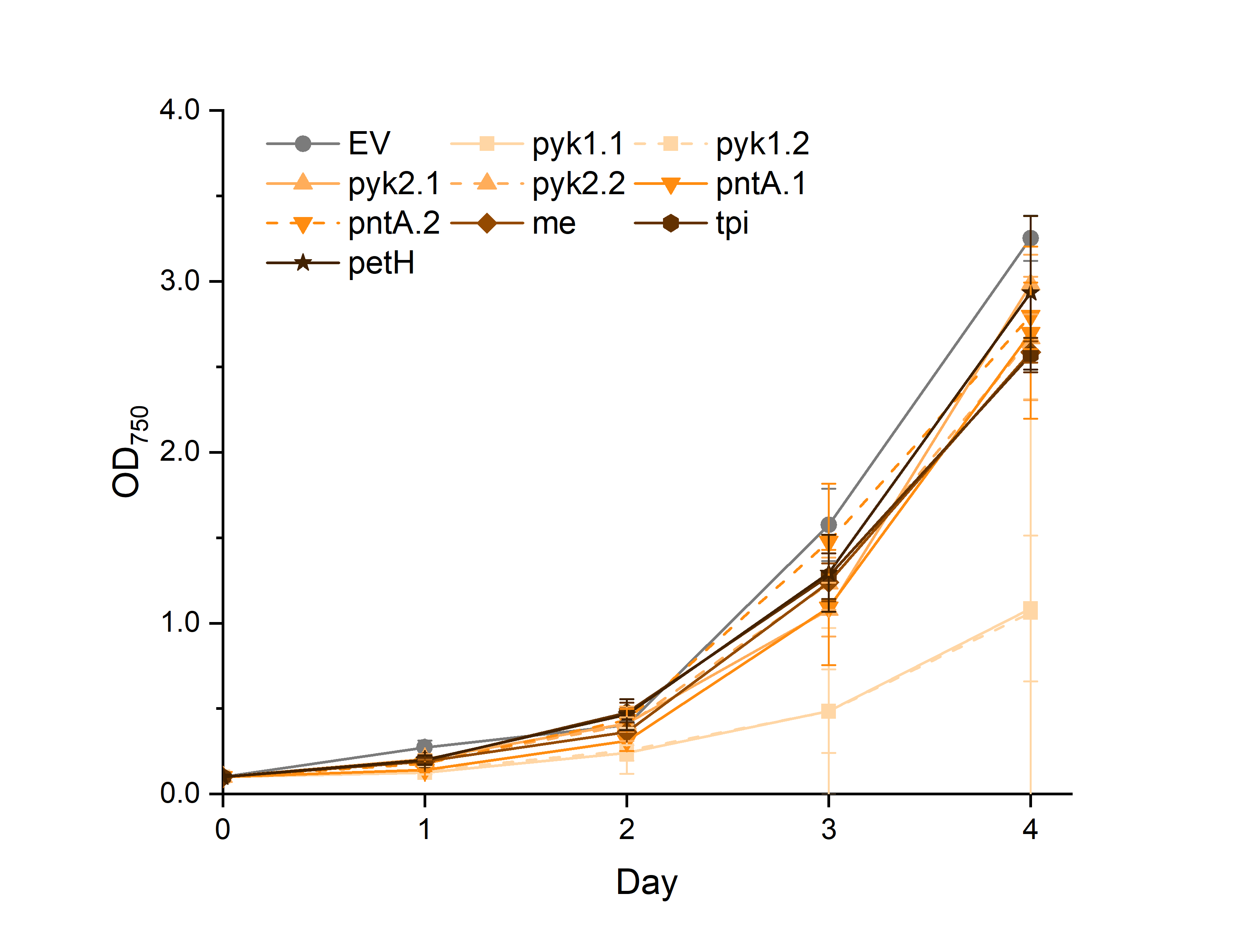

**Supplementary Figure 4.** Growth profiles of strains with CRISPRa targeting for target mapping. EV: negative control. Error bars represent standard deviation (n=3).

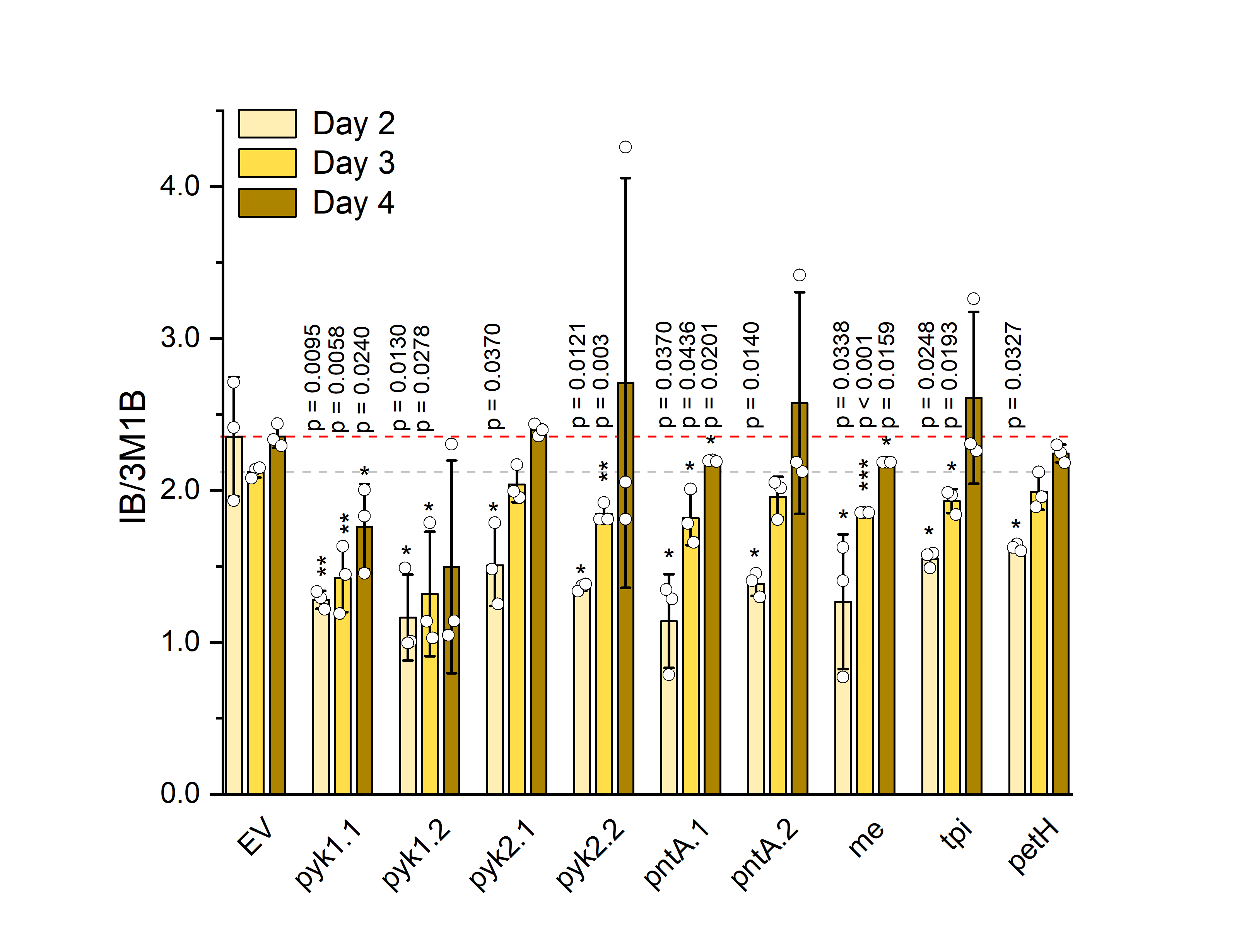

**Supplementary Figure 5.** IB/3M1B ratio for CRISPRa-targeted strains. Error bars indicate standard deviation (n=3). p value representation: * < 0.05; ** < 0.01; *** < 0.001.

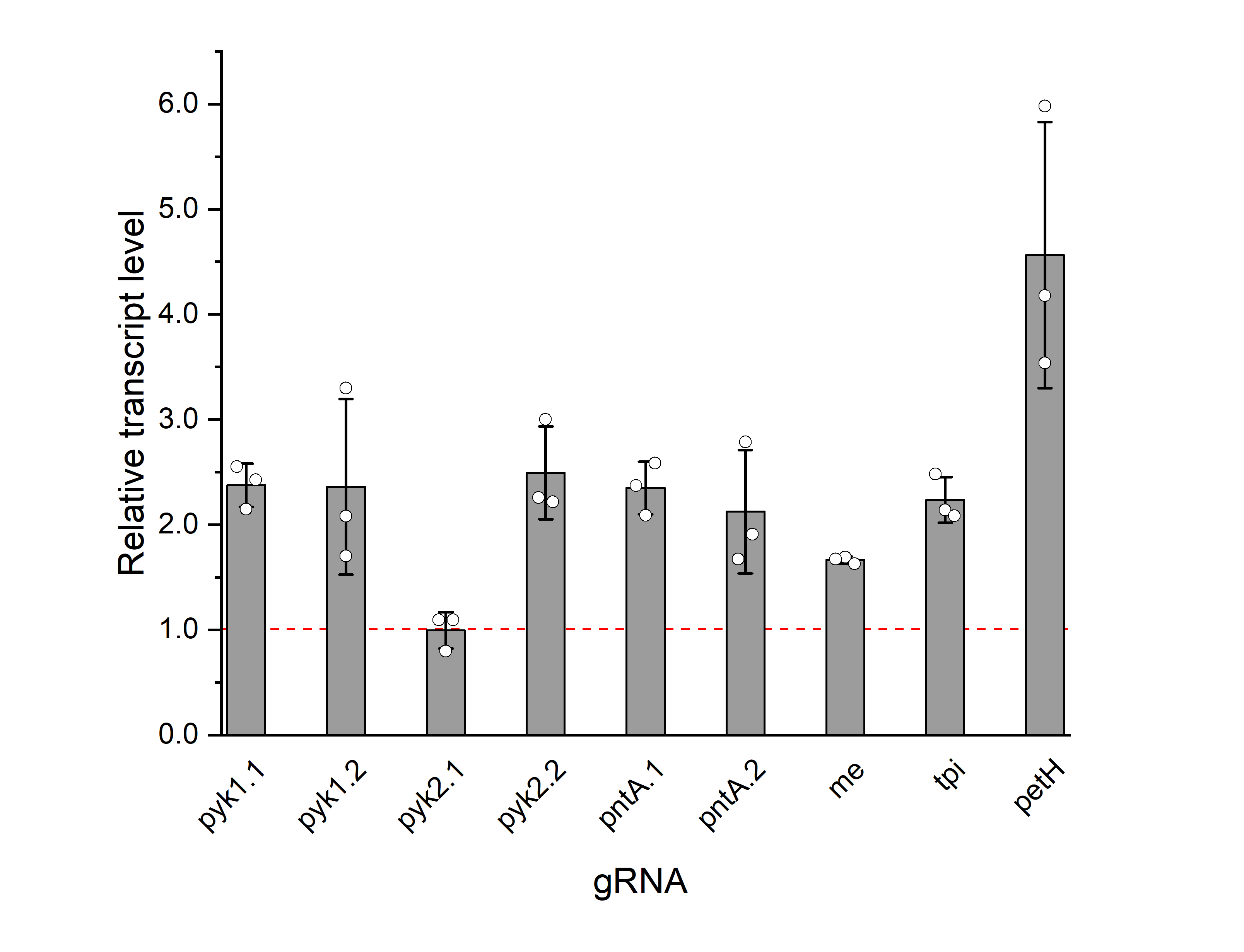

**Supplementary Figure 6.** Relative transcript levels of candidate genes in response to CRISPRa targeting on day 3. Error bars indicate standard deviation (n=3).

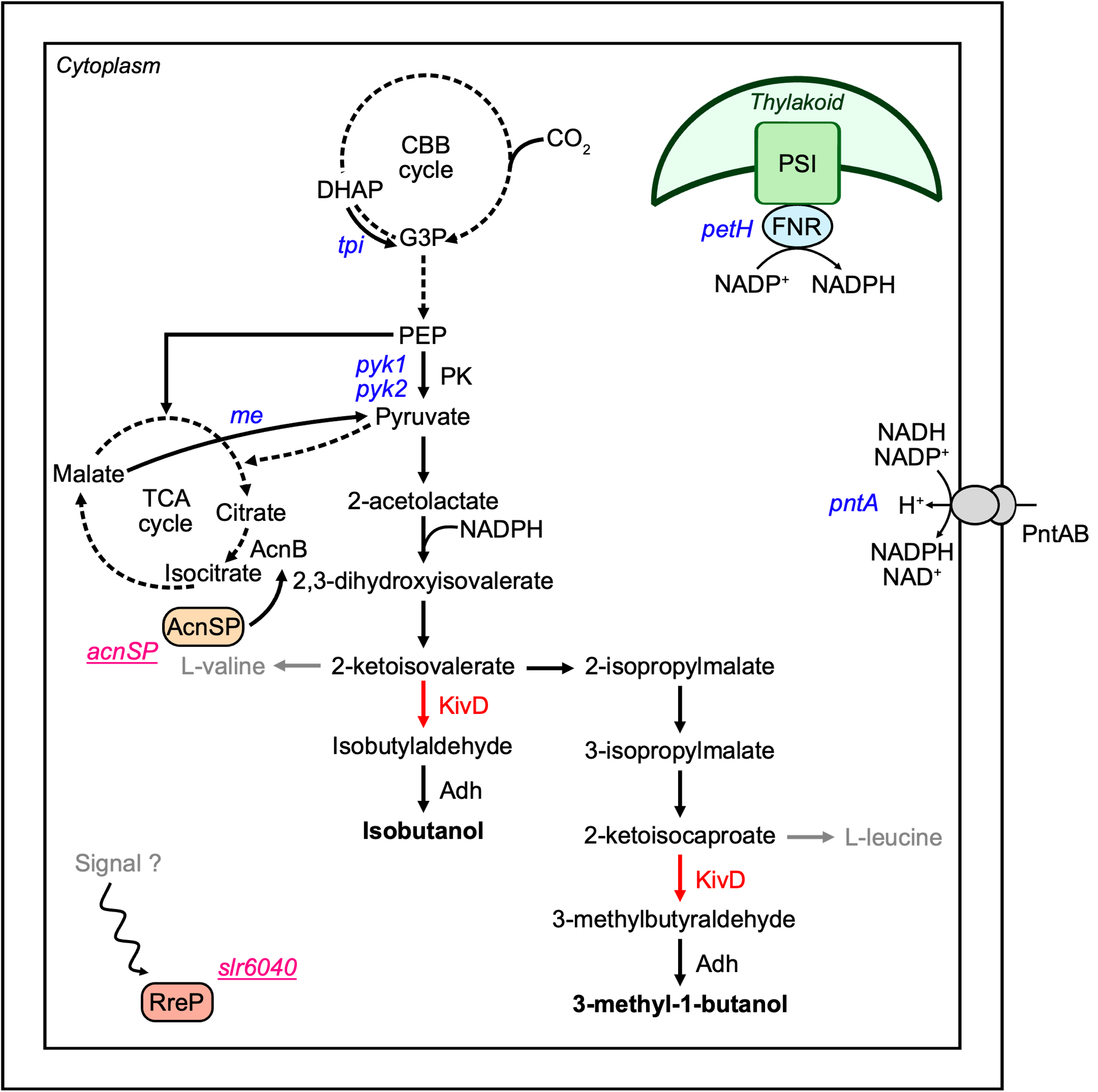

**Supplementary Figure 7.** Simplified metabolic map for IB and 3M1B biosynthesis. Selected genes were upregulated with CRISPRa (blue) to improve pyruvate availability and NADPH regeneration. acnSP and slr6040 (pink and underlined) were targeted with the CRISPRa system for downregulation. Abbreviations: AcnB: aconitase; AcnSP: aconitase small protein; Adh: alcohol dehydrogenase; CBB: Calvin-Benson-Bassham cycle; DHAP: dihydroxyacetone phosphate; FNR: ferredoxin-NADP^+^ oxidoreductase; G3P: glyceraldehyde-3-phosphate; KivD: α-ketoisovalerate decarboxylase; ME: malic enzyme; PEP: phosphoenolpyruvate; PK: pyruvate kinase; PntAB: pyridine nucleotide transhydrogenase; PSI: photosystem I; TCA cycle: tricarboxylic acid cycle.

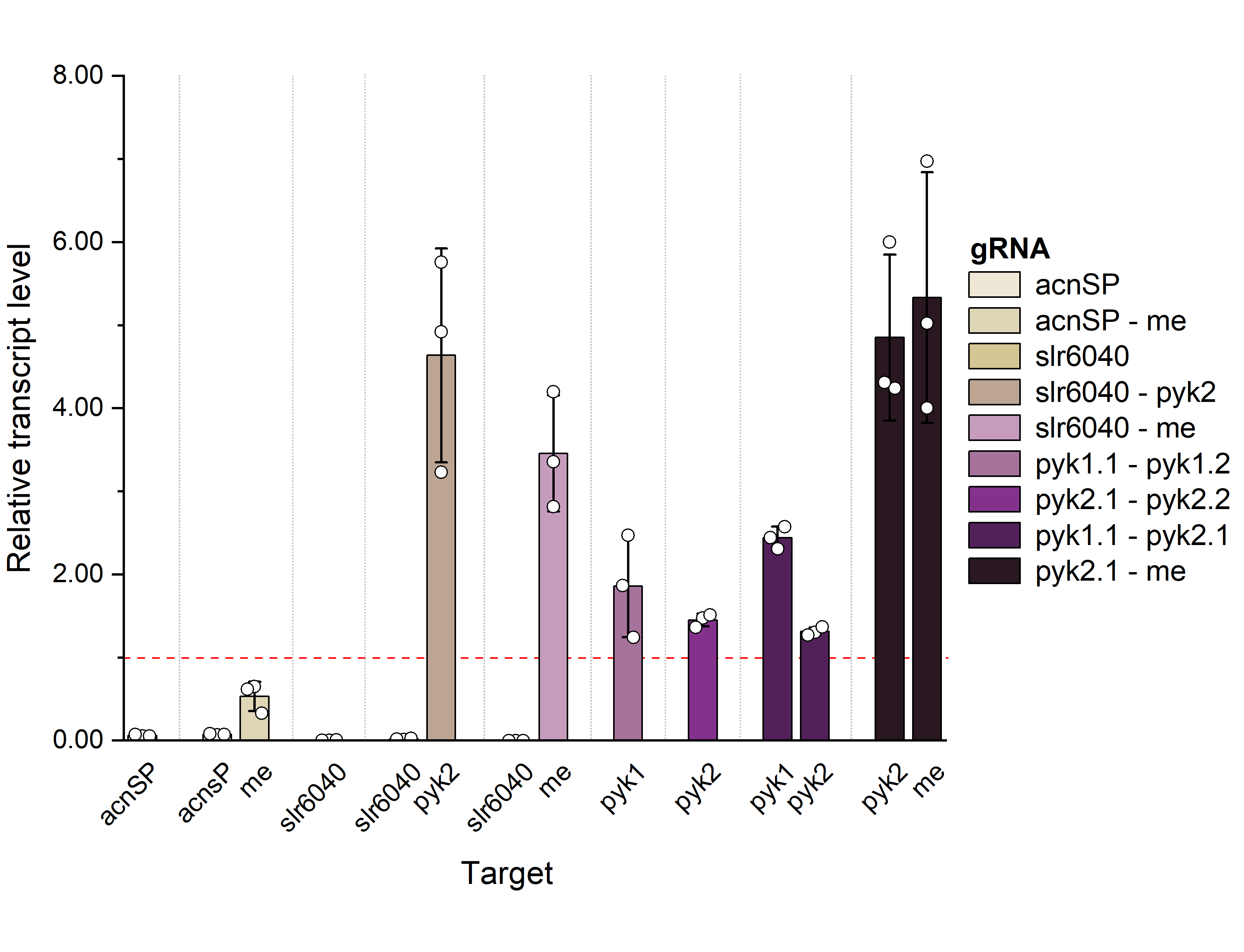

**Supplementary Figure 8.** Relative transcript levels of multiplexed gene targets under CRISPRa-mediated upregulation and repression on day 2. Error bars indicate standard deviation (n=3).
